# Sex-specific developmental changes in spinal cord pain pathways following neonatal inflammation

**DOI:** 10.1101/2023.04.12.536647

**Authors:** Kateleen E Hedley, Annalisa Cuskelly, Rikki K Quinn, Robert J Callister, Deborah M Hodgson, Melissa A Tadros

## Abstract

Early-life inflammation can have long lasting impact on pain processing and pain behaviours. For example, we have shown neonatal inflammation can result in changes within spinal neuronal networks and altered flinching of the hind paw following formalin injection three weeks later. This suggests mechanisms for altered pain behaviours lie in first and second order neurons in the pain neuroaxis. Exactly how these changes progress during postnatal development is not known. Accordingly, we investigated neuroinflammatory markers in sensory neurons (dorsal root ganglia; DRGs) and spinal cords of Wistar rats (both sexes) after early life inflammation. Rats were injected with LPS or saline on postnatal days (P) 3 and 5. DRGs and spinal cords (SC) were isolated on P7, 13 and 21, and the expression of six inflammatory mediators were quantified via RT-qPCR. In the DRG, four proinflammatory mediators were elevated in P7 rats exposed to LPS. By P13, only two proinflammatory agents were elevated, whereas at P21 the levels of all six inflammatory mediators were similar between LPS and saline-treated rats. There were no sex-specific differences in the expression profile of any mediator in DRGs. In the spinal cord this expression profile was reversed with no change in inflammatory mediators at P7, elevation of two at P13 and four at P21 in LPS treated rats. Interestingly, these differences were greater in the spinal cords of female rats, indicating sex-specific modulation of neuroinflammation even at these early stages of postnatal development. The increased inflammatory mediator profile in the spinal cords of P21 LPS-treated rats was accompanied by sex-specific modulation of astrocytic (GFAP) activation, with females showing an increase and males a decrease in GFAP following LPS exposure. Together, these data indicate sensory neurons are more susceptible to acute inflammation whereas inflammation in the spinal cord is delayed. The sex-specific modulation of inflammation during critical phases of development may help explain altered pain behaviours in adult males and females.

## Introduction

Early life insults can drive the expression of chronic disorders which persist well into adulthood. Clinical studies show premature infants exposed to painful stimuli in the NICU have increased responses to heat in early adolescence (Hermann et al., 2006). Similarly, preclinical work in rodent models of early-life pain show rat pups exposed to repeated paw pricks as neonates exhibit exaggerated responses to mechanical stimuli (Anand et al., 1999). Such long-term changes can also be induced by early life inflammation. For example, exposure to lipopolysaccharide (LPS) in the first two postnatal weeks results in enhanced sensitivity to painful stimuli in adulthood (Boisse et al., 2005). Our previous work has validated these alterations in pain perception, with neonatal rats exposed to LPS exhibiting increased pain behaviours in response to formalin injection in adulthood (Zouikr et al., 2015). Most importantly, alterations in formalin-induced pain behaviours emerge one week after neonatal LPS exposure, with postnatal day (P) 13 rats displaying increased licking following formalin injection (Zouikr et al., 2014). This finding was also observed in prenatally-stressed rodents, with P7-8 rats displaying enhanced behavioural responses to formalin injection compared to their non-stressed counterparts (Butkevich et al., 2007). Together, these finding suggest early life inflammation can influence pain perception throughout life.

Although these changes in pain behaviours following neonatal inflammation are well documented, the underlying mechanisms remain elusive. The pain pathway begins in the periphery with the activation of nociceptors located on the peripheral axons of sensory neurons in the dorsal root ganglia (DRG). These signals subsequently enter the spinal cord dorsal horn where second order neurons pass signals along the pain neuroaxis to trigger pain perception and actions (pain behaviours) which reduce pain. Much of the information on neuroimmune responses in the DRG and spinal cord following noxious insults has been obtained in adult animals (see: (Kavelaars and Heijnen, 2021) for review). Few studies have examined the consequences of early life inflammation on first and second order neurons in the early postnatal and preadolescent preclinical models, and even fewer have considered potential sex-specific modulation at these ages. Our previous work showed preadolescent rats exposed to neonatal LPS demonstrated increased flinching following formalin injection at P22 (Zouikr et al., 2014) as well as enhanced neuronal signalling and altered ionic currents in their spinal cord networks (Tadros et al., 2018). Thus, our data examining the pain behaviours of neonatal and preadolescent rats exposed to neonatal LPS implicates the DRG and spinal cord in altering pain behaviours during this early developmental period.

Given the complexity of neuroimmune interactions, upregulation of key elements of this bidirectional communication during critical periods of development is likely to underlie the long-lasting alterations in pain behaviours following neonatal inflammation. However, because there are notable differences in both the nervous and immune systems during these critical periods of development (Zengeler and Lukens, 2021), knowledge gained in adult animals may not apply to neonatal animals. Accordingly, we have examined the profile of key inflammatory markers in both the DRG and spinal cord during the pre-adolescent period following early life exposure to LPS.

## Materials and Methods

### Animals

Experimental animals were derived by breeding naïve male and female Wistar rats (aged 10-12 wks) acquired from the University of Newcastle’s Animal House. Rat pups (n = 40, even split between males and females) were obtained by mating one male with 4 females.

Experiments were undertaken in accordance with federal and state legislation and as approved by the University of Newcastle Animal Care and Ethics Committee. All animals were maintained in a temperature (21°C ± 1) and humidity-controlled environment on a 12 hr light/dark cycle with food (standard rat chow) and water available ad libitum for the duration of the study.

### Intraperitoneal Injection of Lipopolysaccharide

On the day of birth (P1) all pups within a given litter were randomly allocated to receive either lipopolysaccharide (LPS) or saline (SAL) injections. Pups were briefly removed from dams on P3 and given an i.p. injection of either 0.05 mg/kg of LPS (pyrogen free saline and *Salmonella enterica*, serotype Enteritidis) or SAL, and immediately returned to the dams. The injections were repeated on P5 and the pups were then left undisturbed with their dams until experimental timepoints of P7, 13 or 21 to observe any possible changes throughout this developmental period. Animals were sacrificed via guillotine before tissue collection. For each timepoint and tissue type (DRG or lumbar spinal cord) the experimental groups were as follows; female SAL, female LPS, male SAL, male LPS.

### RNA Extraction and RT-qPCR

To quantify inflammatory markers in first and second order components of the pain pathway, thoraco-lumbar dorsal root ganglia (T10 to L3; DRG) and the lumbar spinal cord (T12 – L5) were removed following euthanasia. Both were immediately snap-frozen to -80°C. To extract RNA, separate homogenates from a single DRG per animal and the lumbar region of the spinal cord were obtained. A sample of the resulting homogenate (10-20 mg) was used to assess total cellular RNA using the RNeasy® Mini Kit (Qiagen, Hilden, Germany), according to the manufacturer’s instructions. RNA concentration and quality were measured with a NanoDrop 1000 Spectrophotometer (ThermoFisher Scientific, USA).

Any contaminating genomic DNA within the RNA samples was digested using DNase I (Invitrogen, Scoresby, Australia). Reverse transcription was then performed using Superscript III (Invitrogen, Scoresby, Australia), according to the manufacturer’s instructions. Briefly, 30 – 200 ng of total RNA, 1 μl of oligo(dT)_18_ (Meridian Bioscience, Ohio, USA), 1 μl of random hexar (Meridian Bioscience, Ohio, USA), 1 μl of 10μM dNTP (Meridian Bioscience, Ohio, USA), and molecular grade water to 13 μl, were mixed and heated for 5 min at 65°C in a Thermal Cycler (Eppendorf, Germany). After this step 4 μl of 5x first-strand buffer, 1 μl of 0.1M DTT, 1 μl RNaseOUT (40 U/μl) (Meridian Bioscience, Ohio, USA) and 1 μl SuperScript III RT (200 U/μl) were added and the mixture incubated for 60 min at 50°C, then 70°C for 15 min. Reverse transcription without Superscript III was also undertaken to determine the level of genomic DNA contamination.

All qPCR primers (Table 1) were designed with Ensembl using standard primer design criteria. The primer pairs were then screened through NCBI primer BLAST to ensure primer specificity. A total volume of 12.5 μl was used for the reaction containing; 6.25 μl 2x SensiFAST SYBR (Meridian Bioscience, Ohio, USA), 10 μM each of forward and reverse primers, 1 - 2.5 ng cDNA and molecular grade water to 12.5 μl. After an initial 10 min enzyme activation step at 95°C, 40 cycles at 95°C for 30 s (step 1) followed by 30 s at 60°C (step 2) were completed. Melt curves were generated to confirm the presence of a single PCR product. Primers were deemed specific if a single amplified product was detected by melt curve analysis. Reactions were performed on a 7500 Real Time PCR System (Applied Biosystems, USA) and analysed using the Applied Biosystems 7500 Software (V2.3; Applied Biosystems, USA). For each primer the samples were run in triplicate, including a negative water control on each plate. Delta Ct (ΔCt, threshold cycle) was determined for each gene relative to the housekeeping genes β-Actin and 18S. The ΔΔCt method (Livak and Schmittgen, 2001) was employed for comparisons between groups.

**Table 1.**
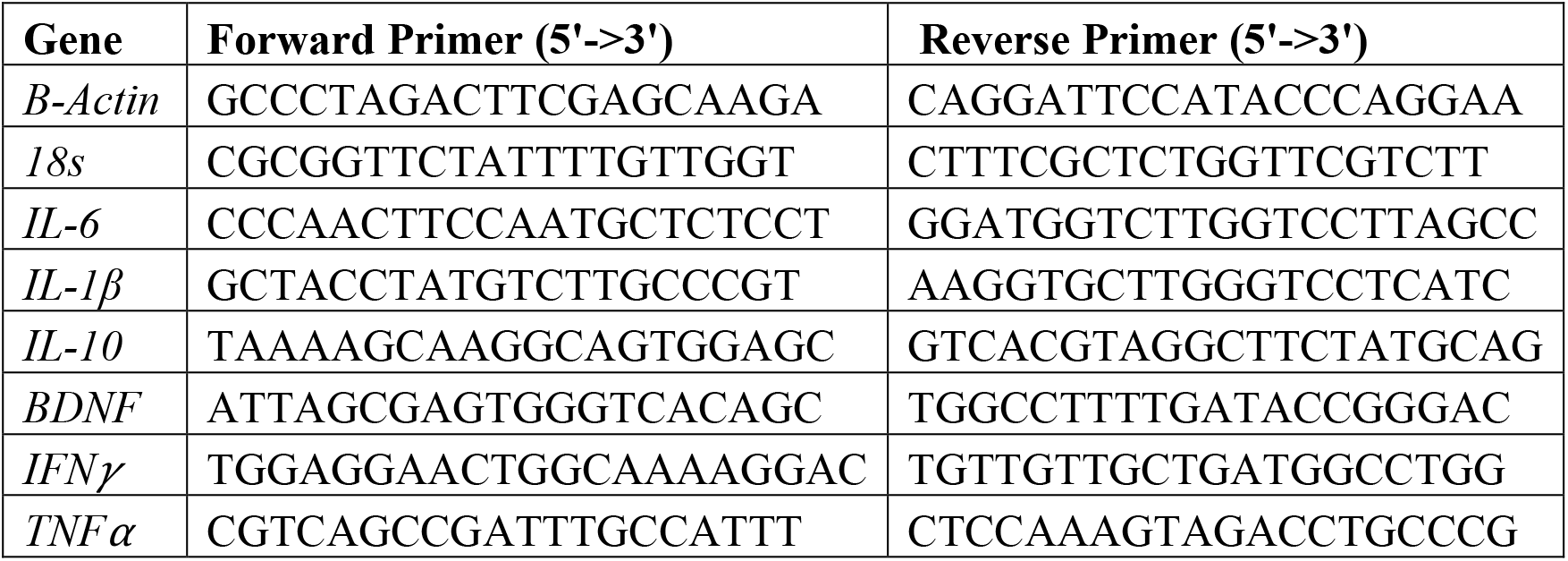
Primer Sequences

### Immunofluorescent labelling of microglia and astrocytes

P21 lumbosacral spinal cords (n = 4 per group) were dissected from euthanised animals and immersion-fixed for 24 hrs in 4% paraformaldehyde (PFA) in phosphate buffer (PBS), then washed in PBS (3 changes). The cords were then washed in 80% ethanol (EtOH), dimethyl sulfoxide (DMSO) and 100% EtOH and embedded in 1450MW polyethylene glycol (PEG). Transverse sections (20 μm thick) were cut on a rotary microtome and collected in PBS. The sections were then blocked in 10% normal donkey serum (NDS, Jackson ImmunoResearch). For labelling with antibodies against ionized calcium binding adaptor molecule 1 (Iba1; for microglia) and glial fibrillary acidic protein (GFAP; for astrocytes), sections were incubated overnight at room temperature in a solution of PBS with 0.1% Triton, 10% NDS, Iba1 (rabbit; 1:250; Wako), and GFAP (chicken; 1:1000; Abcam). After the primary incubation step, sections were rinsed in PBS and incubated for 2 hr at room temperature with secondary antibodies 594-donkey-anti-rabbit (1:50; Abcam) and 488-donkey-anti-chicken (1:50; Jackson ImmunoResearch) and NeuroTrace Blue (1:50, ThermoFisher Scientific). Sections were mounted on slides in buffered glycerol and cover slipped. Images were acquired using an Olympus BX50 microscope equipped with a mercury burner and an Olympus DP72 camera. Mean fluorescent intensity (MFI) was calculated independently for Iba1 and GFAP using ImageJ (National Institute of Health, USA).

### Data analysis and Statistics

Data were analysed via multivariate analysis of variance (ANOVA) using SPSS software (V25; IBM SPSS Statistics, New York, USA). Outlying data points were rejected if they were more than 2 standard deviations from the mean. Post-hoc comparisons were used to determine interaction effects and the Bonferroni correction was used for post-hoc comparisons. All graphs show untransformed means and standard errors of the mean (SEM). Statistical significance was set at p ≤ 0.05.

## Results

### Dorsal root ganglia

Neonatal exposure to LPS altered expression levels of key inflammatory mediators in DRGs (Figures 1-3). These changes were observed at individual time points as well as across the first three weeks of postnatal development. As no sex-specific changes were observed in DRG samples, data for males and females were combined.

**Figure One:**
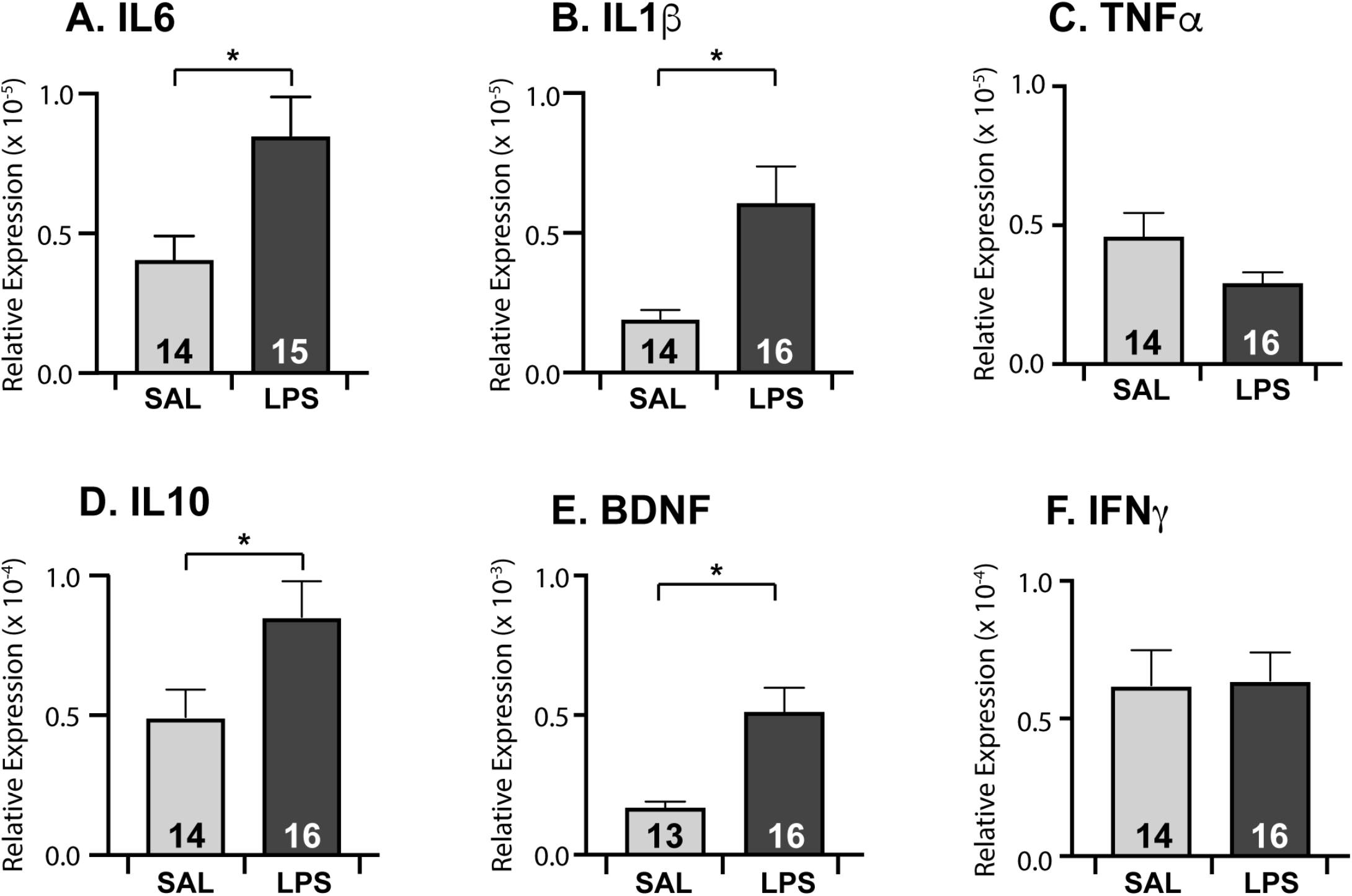
Gene expression levels in dorsal root ganglia at postnatal day 7. Neonatal LPS exposure resulted in an increase in IL6 (A); IL1β (B); IL10 (D) and BDNF (E), with no change in the relative expression of TNFα (E) or IFNγ (F). Data are displayed as mean ± SEM, with the total number of samples shown in each bar. No sex differences were observed, therefore male and female data were combined. * = p < 0.05

**Figure Two:**
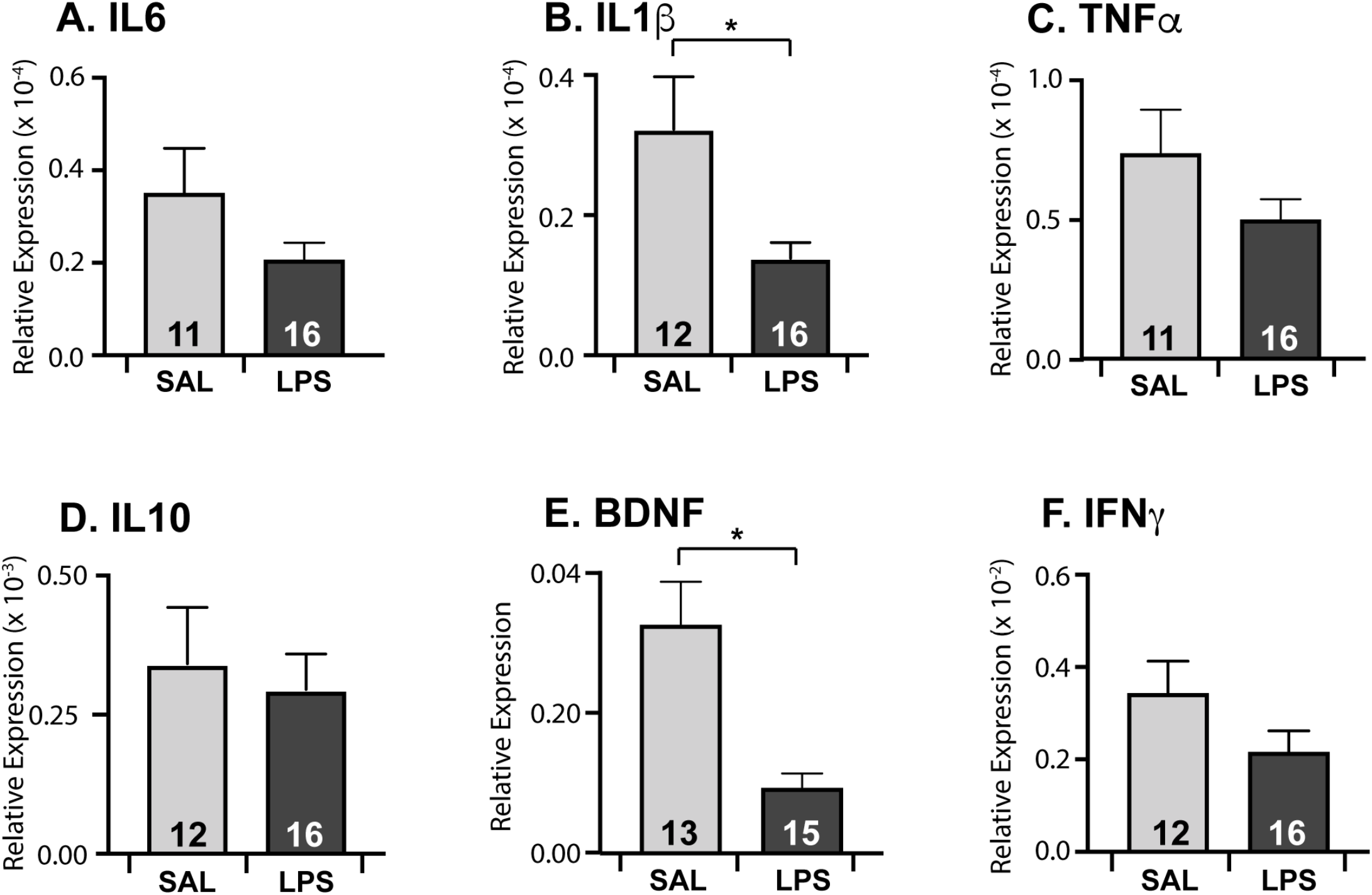
Gene expression levels in dorsal root ganglia at postnatal day 13. Neonatal LPS exposure resulted in a decrease in IL1β (B) and BDNF (E), with no change in the relative expression of IL6 (A); TNFα (C); IL10 (D) or IFNγ (F). Data are displayed as mean ± SEM, with the total number of samples shown in each bar. No sex differences were observed, therefore male and female data were combined. * = p < 0.05

**Figure Three:**
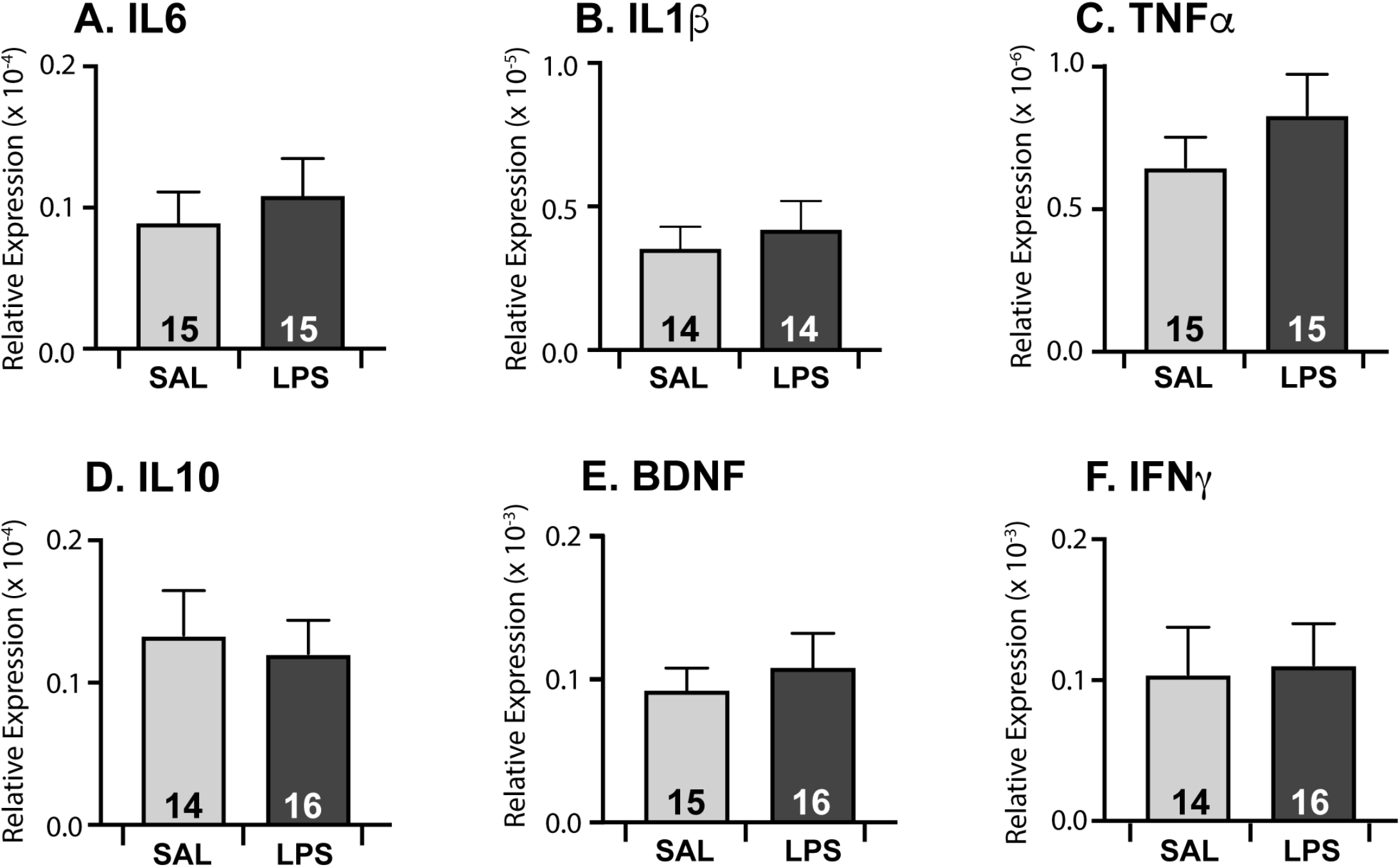
Gene expression levels in dorsal root ganglia at postnatal day 21. Neonatal LPS exposure did not alter gene expression of key inflammatory markers. Data are displayed as mean ± SEM, with the total number of samples shown in each bar. No sex differences were observed, so male and female data were combined.

At P7, expression levels for four inflammatory mediators were increased in the DRGs of LPS treated animals (Figure 1). Specifically, a significant effect of treatment was found for IL6 (F_(1,28)_ = 6.664, p = 0.016), IL1β (F_(1,28)_ = 7.690, p = 0.01); IL10 (F_(1,28)_ = 4.4152, p = 0.045) and BDNF (F_(1,28)_ = 10.047, p = 0.004) with all four markers shown to be elevated in rats exposed to LPS. A similar analysis for the same genes at P13 is shown in Figure 2. At this timepoint only two genes were effected by LPS treatment, and in contrast to P7, their expression levels were lower than their SAL counterparts: IL1β (F_(1,27)_ = 6.892, p = 0.015), and BDNF (F_(1,27)_ = 4.980, p = 0.035). However, neonatal LPS exposure did not affect the expression of any markers in DRGs at P21 (Figure 3). Together, these findings suggest neonatal exposure to LPS induces rapid changes and elevated expression of key inflammatory genes in the DRG. However, these changes in gene expression do not persist past P21.

To assess the impact of LPS on the developmental trajectory of these key inflammatory mediators, we next compared the expression levels for each mediator over the three ages used in this study. In control animals there was a clear peak in gene expression in the DRG at P13 for all mediators: IL1β (F_(1,84)_ = 27.826, p < 0.001); IL6 (F_(1,84)_ = 7.710, p = 0.001); IL10 (F_(1,84)_ = 23.986, p < 0.001); TNFα (F_(1,84)_ = 28.906, p < 0.001); IFNγ (F_(1,84)_ = 50.323, p < 0.001) and BDNF (F_(1,84)_ = 36.017, p < 0.001). LPS exposure interrupted the normal developmental expression of two mediators: IL1β and IL6. Specifically, the peak we observed in IL1β expression in control animals at P13 was not observed in LPS-treated animals, although IL1β expression in both SAL and LPS animals decreased between P13 and P21 (age x treatment interaction: F_(1,84)_ = 8.542, p < 0.001; Post hoc for saline: (F_(2, 78)_ = 30.928, p <0.001; Post hoc for LPS: (F_(2, 78)_ = 3.359, p 0.04). This altered developmental pattern suggests that LPS exposure induces a premature peak in IL1β expression. In addition, whilst saline treated animals displayed an increase in IL6 gene expression between P7 and P13, and subsequent decrease between P13 and P21 (age x treatment interaction: F_(1,84)_ = 3.226, p = 0.045; Post hoc for saline: (F_(2, 78)_ = 10.063, p <0.001), there was no significant effect of LPS treatment on age for IL6. This suggests neonatal LPS exposure flattens the developmental peak of IL6 expression by maintaining elevated levels throughout the first and second postnatal weeks.

### Spinal Cord

Neonatal exposure to LPS resulted in elevated expression of genes for several inflammatory markers in the spinal cord. Like the DRG, these changes occurred at specific time points during the first three weeks of postnatal development. However, in contrast to the DRG, sex-specific changes in gene expression were observed. Accordingly, gene expression data are presented separately for male and female animals (Figures 4-6).

**Figure Four:**
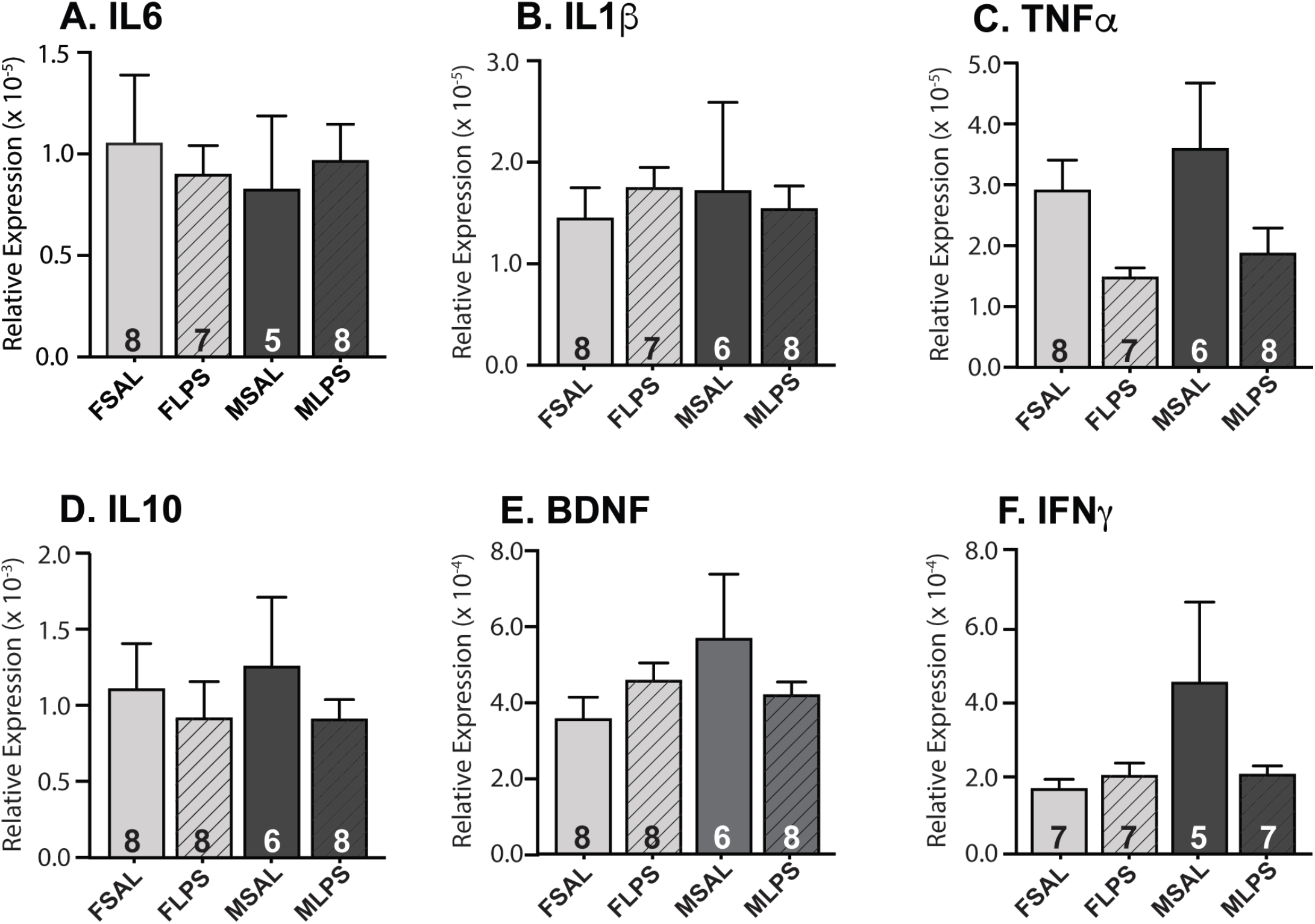
Gene expression levels in spinal cord at postnatal day 7. Neonatal LPS exposure did not result in any changes in gene expression of key inflammatory markers. Data are displayed as mean + SEM, with the total number of samples shown in each bar.

**Figure Five:**
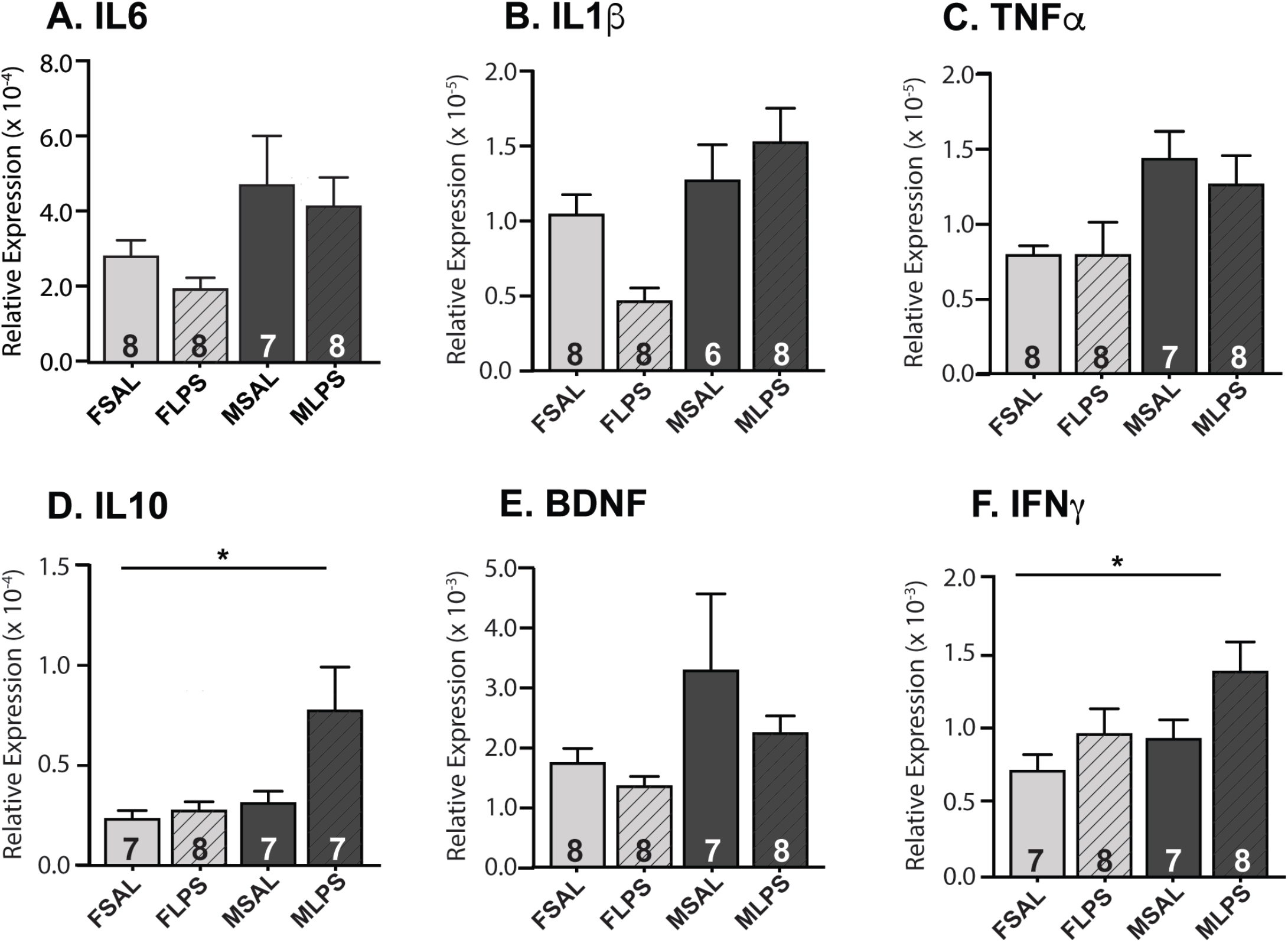
Gene expression levels in spinal cord at postnatal day 13. Neonatal LPS exposure resulted in a significant increase IL10 (D) and IFNγ (F) gene expression but no change in IL6 (A), IL1β (B), TNFα (C) or BDNF (E). Data are displayed as mean ± SEM, with the total number of samples in each bar. * = p < 0.05

**Figure Six:**
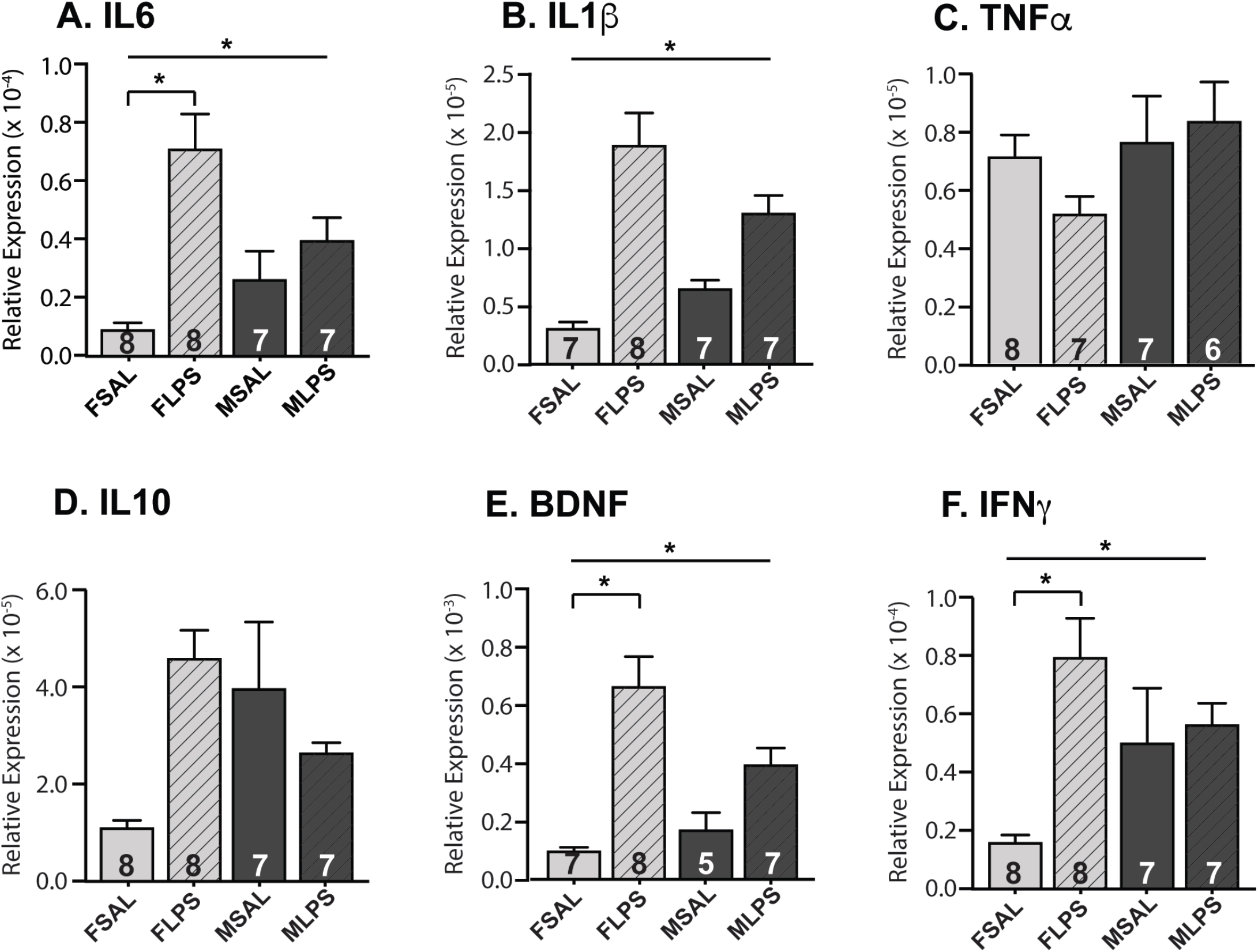
Gene expression levels in spinal cord at postnatal day 21. Neonatal LPS exposure resulted in a significant increase IL1β (B) gene expression. Females showed significant increases in gene expression of IL6 (A), BDNF (E) and IFNγ (F), with no changes in gene expression for males. No changes were observed in TNFα (C) or IL10 (D). Data are displayed as mean ± SEM, with the total number of samples in each bar. * = p < 0.05

There was no effect of LPS on gene expression for any marker at P7 in both sexes (Figure 4). However, by P13, expression levels of two inflammatory mediators were impacted by neonatal LPS, and this occurred in a sex-specific manner. Specifically, IL10 (F_(1,26)_ = 4.7586, p = 0.0391) increased in LPS treated males compared to their male and female counterparts (Figure 5). IFNγ expression levels also increased in males but only relative to females (F_(1,26)_ = 6.854, p = 0.015). At P21, neonatal LPS exposure induced greater and more complex changes in the gene expression levels of four key inflammatory mediators (Figure 6). LPS treated animals exhibited increased expression of IL6 (F_(1,27)_ = 20.808, p < 0.001), IL1β (F_(1,27)_ = 33.692, p < 0.001), BDNF (F_(1,27)_ = 12.141, p = 0.002) and IFNγ (F_(1,27)_ = 8.819, p = 0.006). These increases differed in males and females, with females showing increased expression of IL6, IFNγ and BDNF, which were not observed in males (IL6 - treatment x sex interaction: F_(1,27)_ = 7.935, p = 0.009; Post hoc for females: F_(1,25)_ = 27.622, p = <0.001; IFNγ - treatment x sex interaction: F_(1,27)_ = 5.9181, p = 0.022; Post hoc for females: F_(1,25)_ = 15.635, p = 0.001; BDNF - treatment x sex interaction: F_(1,27)_ = 12.141, p = 0.002; Post hoc for females: F_(1,25)_ = 19.136, p = <0.001). Together, these data suggest LPS exposure does not immediately affect inflammatory mediator gene expression in either sex as it does in DRG. Rather, female rats exposed to LPS exhibit a delayed increase in key inflammatory markers two weeks after the initial inflammatory insult.

To further explore the impact of LPS exposure on the developmental trajectory of these key inflammatory mediators, we next compared the expression levels for each mediator across the three ages. In contrast to the DRG, only three of the six mediators demonstrated a clear peak in gene expression at P13 in control animals (IL6 (F_(2,84)_ = 53.648, p = <0.001); IFNγ (F_(2,84)_ = 83.229, p < 0.001) and BDNF (F_(2,84)_ = 25.913, p < 0.001)). Expression of the remaining mediators, IL10, TNFα and IL1β decreased from P7 to P21 without a discernible peak in the second postnatal week (IL10 (F_(2,84)_ = 57.406, p < 0.001); TNFα (F_(2,84)_ = 8.508, p < 0.001); IL1β (F(2,84) = 3.056, p = 0.05). The expression of IL1β and IFNγ were both developmentally modulated by neonatal LPS exposure, with levels of both mediators remaining high at P21 despite decreasing compared to P13 in saline-treated rats (IL1β: age x treatment interaction: F_(2,84)_ = 4.474, p = 0.014; Post hoc for LPS: F_(2, 78)_ = 3.684, p = 0.03; Post hoc for saline: F_(2,,84)_ = 4.028, p = 0.022; IFNγ: age x treatment interaction: F_(2,,84)_ = 4.555, p = 0.013; Post hoc for LPS: F_(2, 78)_ = 65.606, p = <0.001; Post hoc for saline: : F_(2, 78)_ = 24.806, p <0.001). Together, this suggests LPS disrupts the normal developmental trajectory of key inflammatory mediators the spinal cord, by maintaining elevated expression of IL1β and IFNγ into the third postnatal week.

### Spinal microglia and astrocytes

Because the increase in inflammatory mediators in the spinal cord was greatest at P21 (Figure 6), we next examined microglia and astrocyte modulation after LPS exposure in the pain processing region of the spinal cord (ie, the superficial dorsal horn, SDH; Figure 7). There was no difference in the mean fluorescence intensity for Iba1 in any group (Figure 7E). In contrast, changes were observed in GFAP fluorescent intensity and the nature of the change differed between sexes (sex x treatment interaction F_(1,14)_ = 12.892, p = 0.04). GFAP mean fluorescent intensity increased in females after neonatal LPS exposure (Post hoc for females F_(1,11)_ = 5.071, p = 0.046), whereas it decreased in males (Post hoc for males F_(1,11)_ = 8.138, p = 0.016). These data suggest neonatal exposure to LPS had no impact on microglial activation in the SDH at P21 in either male or female rats. In contrast, astrocyte activation is modulated in a sex-specific manner in the SDH at P21 following neonatal LPS exposure.

**Figure Seven:**
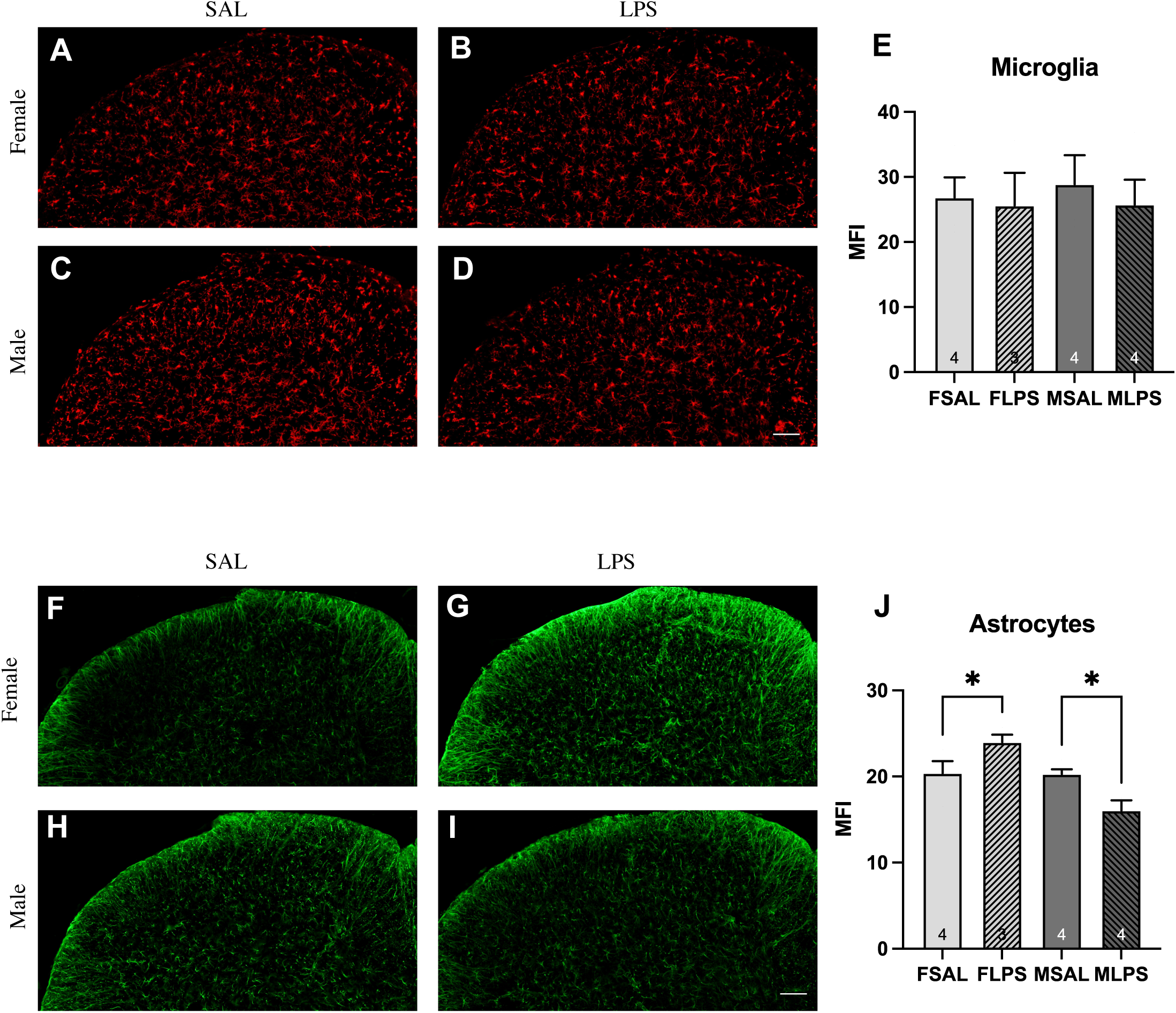
Microglia and astrocyte immunofluorescence in the superficial dorsal horn of the spinal cord at P21. Mean fluorescent intensity (MFI) of Iba1 immunofluorescence (A-D) did not differ between the groups examined (E). In contrast, females exposed to neonatal LPS (G) showed significantly higher GFAP MFI (J) compared to saline-injected controls (F), whilst males exposed to neonatal LPS (I) showed a significantly lower MFI (J) compared to their saline counterparts. Data are displayed as mean ± SEM, with the total number of samples shown in each bar. * = p < 0.05

## Discussion

Our study demonstrates an acute upregulation of key inflammatory mediators in DRGs immediately following LPS exposure, which resolves within two weeks of the neonatal insult. This is followed by an increase in the same suite of inflammatory mediators in the spinal cord in pre-adolescent animals. Interestingly, these differences in the spinal cord are sex-specific, with females showing a greater increase in gene expression of inflammatory mediators than males, even though these animals are pre-adolescent. Furthermore, these sex-specific differences are matched by sex-specific modulation of the immunoreactivity of astrocytes in the superficial dorsal horn, with females showing increases whilst males exhibit decreased GFAP levels. These data emphasise the need for sex-specific studies of neuroimmune interactions throughout the developmental period, at all nodes of the pain pathway in order to elucidate the mechanisms underlying the long-term impacts of neonatal immune activation.

### Evidence for an indirect impact on spinal cord networks

We observed significant interactions between age and treatment in both the DRG and spinal cord, suggesting neonatal LPS exposure interrupts the normal developmental trajectory of these two critical regions in the pain neuroaxis. For the DRG, the inflammation observed at P7 appeared to resolve soon after LPS exposure, with no changes in inflammatory profile at P21. In contrast, we did not observe changes in the P7 spinal cord, but found significant evidence of increased inflammation at P21 – ie, two weeks after the initial LPS insult. This suggests a direct impact of LPS on peripheral DRG, but an indirect mechanism for alterations in spinal networks. Moreover, these interactions were also observed in a sex-specific manner in the spinal cord, with females displaying greater deviation from the saline controls than males.

Intriguingly, this is in contrast to prior studies on neonatal LPS, with prior studies reporting increases in the expression of key inflammatory mediators spinal cord within 24 hours of an injection of LPS at similar developmental timepoints (Hunter et al., 2014;Hsieh et al., 2018). One of the key findings of the present paper is an augmented response in the spinal cords of female rats at P21. Whilst Hsieh et al did compare the sexes and found no difference between males and females in their study, Hunter et al do not report any sex-specific considerations at all. Therefore, the sex-specific modulation we observed may have been obscured by the combination of both sexes in the prior studies. In addition, both prior studies utilised only a single injection of LPS at the desired timepoint and examined cytokine levels within 24 hours of exposure, whereas we utilised two injections across at P3 and P5 and documented modulation of cytokine levels two weeks after the initial insult. It is well known that the exact doses, timing and ages of LPS exposure influence the final outcome (Bao et al., 2022), subsequently it is hard to make a direct comparison between different applications of LPS during this critical window of development.

Cytokines are known to be essential in neurodevelopment (see (Zengeler and Lukens, 2021) for review), with many interleukins and grow factors implicated in neurogenesis, differentiation and even the formation and maintenance of synapses. Moreover ‘critical periods’ during development exist for the above processes. Thus the role these cytokines play in neurodevelopment is complex, and highly dependent upon the developmental stage, microenvironment and activation of signalling pathways. The modulation of key inflammatory mediators observed in this study occurred over known critical periods in spinal cord development (Walsh et al., 2009) during which expression of certain types of potassium channels that contribute to neuronal excitability change markedly. These potassium channels are known to be modulated by elevated cytokines arising from maternal immune activation in CA1 hippocampal neurons (Griego et al., 2022), suggesting a complex relationship exists between the altered cytokine levels and ongoing neurodevelopment following neonatal inflammation.

### The role of glia in the long-term impact of early life inflammation

Interestingly, we observed no change in Iba1 immunofluorescence in either sex at P21, suggesting microglia are not involved in the altered levels of inflammatory mediators we observed in the female spinal cord. This is surprising, given microglia in the brain have been shown to react strongly to early life stress (for review see: (Schwarz and Bilbo, 2012).

However, astrocytes can also be influenced by peripheral inflammation to produce a suite of inflammatory mediators, which can be either pro-inflammatory and neurodestructive, or anti-inflammatory and neuroprotective (see: (Colombo and Farina, 2016) for review). In our sample, we observed a concurrent increase in both pro inflammatory cytokines IL6 and IL1β and anti inflammatory mediators IFNγ and BDNF in the spinal cord of P21 females exposed to neonatal LPS, as well as an increase in GFAP immunofluorescence. In contrast, there were no differences in these inflammatory mediators in male P21 spinal cords, and GFAP immunofluorescence decreased. Therefore, it is plausible that female spinal astrocytes are modulated by early life LPS exposure and are implicated in the altered levels of inflammatory mediators. More work however is needed to determine whether this modulation is neuroprotective or destructive.

In naïve animals, male cortical astrocytes reach a ‘mature’ phenotype at P7, whilst female astrocytes reach the same level of maturity later at P14 (Rurak et al., 2022). While sex-specific alterations of glia within the brain are known to occur in a number of laboratory models of early life inflammation (for review see: (Schwarz and Bilbo, 2012), few studies have examined the impact on glia within the spinal cord. Spared nerve injury on male rats during early postnatal development (P0-P21) impacts spinal cord astrocytes more heavily than microglia (Vega-Avelaira et al., 2007). Moreover, this study also showed upregulation of GFAP was delayed in younger animals - changes were not observed until 5 days after an injury inflicted during the first postnatal week. Similarly, the impact of neonatal anoxia on sensorimotor processing demonstrated increased GFAP, as measured by western blot, in the spinal cord of adult female rats exposed to neonatal anoxia. Importantly, this aligned with a decrease in mechanical and thermal thresholds for paw withdrawal (Helou et al., 2021). Interestingly, although adult males exposed to neonatal anoxia also displayed decreased mechanical thresholds, there was no change in their spinal cord GFAP levels. Together with the findings of this study, these data suggest astrocytes are more likely to show long term modulation following neonatal inflammation, and this modulation occurs in a sex-specific manner.

### Implications for pain behaviours

Inflammatory mediators are known to have a direct impact on neuronal signalling, with many cytokines exerting an influence on specific elements of neuronal communication. Within spinal cord networks, IL6 reduces inhibitory signalling and BDNF enhances excitatory signalling (Lu et al., 2007)). Furthermore, IL1β causes concurrent increases in excitatory and decreases in inhibitory signalling (Gustafson-Vickers et al., 2008) and an imbalance and overall increase in excitation of spinal cord networks. This is important because enhanced spinal cord excitation would increase signalling along the entire pain pathway, ultimately enhancing the signal reaching the brain and impacting pain perception and behaviours. At P21, we observed a significant increase in IL6, IL1β and BDNF in the spinal cord of females exposed to neonatal LPS, suggesting enhanced signalling exists in the spinal neuronal network these animals. Indeed, our previous work show increased excitatory currents within spinal cord networks at P22 (Tadros et al., 2018), as well as heightened flinching in P22 animals following neonatal LPS (Zouikr et al., 2014). Together, our work on the spinal cord of preadolescent rats exposed to neonatal LPS suggest the neural processing within spinal cord networks plays a role in altering pain behaviours following early life inflammation.

Gender-differences in clinical pain conditions are widely described in the literature (see (Mogil, 2012)for review), however, the evidence for sex-specific modulation of spinal cord neural processing is less clear. Research examining the impact of neonatal hind paw incision, a laboratory model that mimics neonatal surgery, demonstrated a reduction in inhibitory signalling within neural networks of adult spinal cords exposed to the neonatal incision. However, there were no differences between the modulation observed in male and female adults (Li and Baccei, 2019). Follow up work in adult female mice exposed to the neonatal hind paw incision also demonstrate an imbalance in the excitatory/inhibitory connections of a subpopulation of spinal cord neurons (Brewer et al., 2020), however similar experiments were not carried out in male animals, so it is unclear whether this is particular imbalance is sex-specific. Clearly, spinal cord networks and their associated processing of nociceptive signals is susceptible to early life interventions, and the data in the present study highlights the need to examine changes at multiple stages of development, not just in adulthood.

Together, our findings provide evidence of an indirect impact of neonatal LPS on spinal cord networks, with sex-specific modulation occurring prior to adolescence. We found evidence of altered inflammation in the spinal cord at P21, which was the latest age examined in this study. Additionally, these changes did not emerge until two weeks after the initial LPS insult, implicating an indirect mechanism in this modulation of inflammatory mediators. Alongside these altered inflammatory mediators, we also observed sex-specific modulation of astrocytes at P21, which could contribute to activation of both neurodestructive and neuroprotective pathways. This corroborates the concept of “priming”, whereby early life insults alone may not result in an overtly altered phenotype, however, it may increase the impact of a second insult later in life. This underlies the development of two-hit laboratory models which demonstrate an exaggerated response in adult animals pre-exposed to LPS during early development (for examples: (Zhao et al., 2008;Tufvesson-Alm et al., 2020). This highlights the importance of examining both sexes during the early postnatal developmental period to fully elucidate the role of early life events in shaping future pain behaviours.

